# Convergent adaptation in mitochondria of phylogenetically distant birds: does it exist?

**DOI:** 10.1101/2020.09.15.298117

**Authors:** Valentina Burskaia, Ilja Artyushin, Nadezhda Potapova, Kirill Konovalov, Georgii A. Bazykin

## Abstract

In a wide range of taxa, proteins encoded by mitochondrial genomes are involved in adaptation to lifestyle that requires oxygen starvation or elevation of metabolism rate. It remains poorly understood to what extent adaptation to similar conditions is associated with parallel changes in these proteins. We search for genetic signal of parallel or convergent evolution in recurrent molecular adaptation to high altitude, migration, diving, wintering, unusual flight abilities, or loss of flight in mitochondrial genomes of birds. Developing on previous work, we design an approach for detection of recurrent coincident changes in genotype and phenotype, indicative of an association between the two. We describe a number of candidate sites involved in recurrent adaptation in *ND* genes. However, we find that the majority of convergence events can be explained by random coincidences without invoking adaptation.

## Introduction

Mitochondrial genes were repeatedly claimed to adapt in response to lifestyles that require oxygen starvation or elevation of metabolism rate in a wide range of eukaryotic taxa (Das 2006). The most obvious candidate species for evolution of hypoxia tolerance are those inhabiting high altitudes. Indeed, high-altitude adaptations have been described for mitochondrial genes of many species, including the COX3 gene of the bar-headed goose (Scott 2011), ATP6 and ATP8 in shrimps from genus *Artemia* (Zhang, 2013), ND5 in caterpillars of genus *Gynaephora* (Yuana et al 2018), COX1 in Tibetan antelope *Pantholops hodgsonii* (Xu et al, 2005), and ND2, ND4, and ATP6 in Tibetan galliform birds (Zhou, 2014). Adaptations to high altitude in human populations typically involve mutations in ND1 as well as probably other genes (Kang et al, 2013, Ji et al, 2012). Besides high-altitude hypoxia, some rodents likely developed adaptations to subterranean hypoxia (Tomasco, 2011).

Life at extreme temperatures and extraordinary physical activity could also cause adaptive evolution in mitochondrial ge nes through their effect on energy metabolism. It has been shown that genes involved in oxidative phosphorylation could take part in adaptation to arctic environment: adaptations in *ND1, ND3* and *ND4* genes were described in the Atlantic salmon (Consuegra et al, 2015), and adaptations in *CYTB* gene were found in European anchovy (Silva et al 2014). Furthermore, there is evidence for adaptation to long-range migrations in the yellow-rumped warbler (Toews, 2014). Flight is another energy-consuming adaptation, and some studies confirm adaptation of mitochondrial genes to flight in bats (Shen et al 2010).

Most evidence for selection in mitochondria is indirect. It often comes down to description of differences in frequencies of a few alleles, which could be a consequence of random drift in small populations. Positions discovered in different studies rarely overlap, suggesting that different organisms use different mechanisms of OXPHOS system adaptation to similar environments, or that some of the findings are erroneous. Still, some gene regions are more likely to be affected by adaptation (da Fonseca et al, 2008). Furthermore, the role of mitochondrial electron transport chain in physiological acclimation was demonstrated in many experimental studies. For example, gene expression, protein abundance, and the enzyme activity changes in plants and animals in process of cold acclimation (Armstrong et al 2008, Lucassen 2006).

To study this systematically, we decided to conduct a broad search for adaptive convergent evolution in mitochondria of birds in an attempt to find universal genotype-to-phenotype associations. We concentrated on mitochondrial adaptations in bird species that likely face hypoxia (high altitude and diving) or requirement for elevated (long-distance migration, wintering at high latitudes and unusual flight abilities) or reduced (loss of flight) rate of metabolism, hypothesizing that detected adaptations may confer resistance to these types of physiology.

To estimate potential convergence, we develop upon an existing phylogenetic test for detection of parallel adaptation (TreeWAS package, Collins and Didelot, 2018). This test is based on reconstruction of ancestral states in the internal nodes of the phylogeny, and then counting the number of coincident changes of phenotype and genotype at each amino acid site at the same branch of the phylogenetic tree. Additionally, at each site, we measure the change of amino acid propensities associated with phenotype change (PCOC package, Rey et al, 2018). While we detect a number of candidate sites that could be associated with convergent adaptation to high altitude and long-distance migration in birds, we only detect one site that was associated according to both tests, and this association was borderline significant. Overall, we find little or no signal of recurrent adaptation, indicating that adaptation to extreme physiology in birds can proceed via different routes in different species, and/or that it can be largely driven by non-mitochondrially encoded genes.

## Materials and Methods

### Phenotypes

We analyzed 415 species of birds. All species were divided into 7 groups according to their phenotypic characteristics. Phenotypes were classified in accordance with the Bird of the World research database (https://birdsoftheworld.org, accessed August 20, 2020). As high altitude we classified those species for which the lower boundary of the range was over 2000 meters, and the upper boundary of the range, over 4000 meters above sea level. As divers we classified those species which can spend at least several minutes underwater. As species with ability for long-distance migration we considered those species with non-overlapping or weakly overlapping breeding and wintering ranges. As wintering we classified those species which are typically exposed to sub-zero temperatures and snow cover for many months each year. We also formed two samples of species with specific flight-related phenotypes: flightless birds, a phenotype which has originated repeatedly in different groups of island birds; and birds with outstanding flight abilities. The latter group united swifts, hummingbirds, swallows, terns and gulls, scuas, gannets, tropicbirds, falcons and accipiters. Although the similarity of these adaptations may be controversial, we hypothesize that the lifestyles of all these groups involve high energy demand and thus could affect the mitochondrial genes similarly.

To study phenotypic associations, we also need a reference group of species which do not carry the specific adaptations considered in this work. The choice of such a reference is a complicated task, because of the complexity of natural ecological adaptations. As the reference group, we decided to use tropical and subtropical birds with ranges not extending above 2500 meters and which have none of the specific aforementioned adaptations. The number of species in each phenotypic group is provided in Table 1, and the list of species is provided in Figure S5.

**Table 1:** Number of species in each phenotypic group.

|  | Number of species | Number of times the phenotype emerged independently |
| --- | --- | --- |
| High altitude | 23 | 7 |
| Diving | 25 | 3 |
| Long distance migration | 91 | 33 |
| Wintering | 28 | 11 |
| Flightless | 33 | 6 |
| Outstanding flight abilities | 58 | 7 |
| Reference | 174 | - |

### Gene sequences and phylogeny

We downloaded complete mitochondrial sequences of birds from Genbank (https://www.ncbi.nlm.nih.gov/genbank/, accessed August 20, 2020). The sequence of each of the 13 genes was obtained according to the GenBank annotation. Species with duplicated genes were excluded from analysis. Sequences of each gene were aligned independently with MACSE toolkit (Ranwez et al, 2011). Columns of alignment with gaps were excluded by trimAl software (Capella-Gutierrez et al, 2009). Phylogenetic analysis was performed in IQ-Tree 2 package (Minh et al, 2020) based on nucleotide alignment, split into 39 partitions by gene and codon position. As early divergences in the bird tree of life are discordant (Jarvis et al, 2014), we used constraints for bird orders branching in our reconstruction. We applied constraints from the most recent revision of bird phylogeny (Kimball, 2019), which combines nuclear and mitochondrial data to construct a consensus supertree for 707 bird species. Amino acid ancestor states reconstruction was also performed in IQ-Tree with usage of genewise partitioned mtVer evolutionary model.

### Search for convergent evolution and phenotype to genotype associations

To count simultaneous changes of phenotype and genotype at the site, we use the simultaneous score of the TreeWAS package (Collins and Didelot, 2018). The simultaneous score is designed for the so-called phylogenetic GWAS analysis. It splits each alignment column into binary (two-state) SNPs and counts simultaneous changes of phenotype and genotype at tree branches. To estimate the probability that the observed association is non-random, TreeWAS simulates a “null” genetic dataset under the empirical phylogenetic tree and terminal phenotypes. It also takes from empirical data the distribution of numbers of substitutions per site. In each simulation, it counts the number of phylogenetic branches with simultaneous changes of phenotype and genotype, and combines these counts to obtain the null distribution. At the upper tail of the null distribution, a threshold of significance is drawn at the quantile corresponding to [1 - (alpha-level corrected for multiple testing)]. If a locus in the real dataset has more simultaneous changes than the threshold, it is considered to be significantly associated with the corresponding phenotype.

TreeWAS was originally developed for analysis of whole-genome datasets of closely related species, in which the assumption that no more than two variants may occur in any particular site generally holds. By contrast, in distantly related bird species considered here, the amino acid sequence of mitochondria has frequently undergone multiple substitutions per site. Therefore, we had to adapt TreeWAS for dealing with non-binary SNPs. This is non-trivial, as we have no a priori knowledge which of the amino acid changes may be associated with changes in the phenotype.

We used three approaches to identify the phenotype and genotype changes. Generally, each approach resulted in its own set of branches with phenotype and genotype changes, and therefore in different estimates of the simultaneous score.

In the first and in the second approach, we designated one amino acid as foreground, and did not distinguish between the remaining amino acids. We also designated one phenotype (of the two possible phenotypes) as foreground. In the first approach, as genotype changes, we only considered the gains of the foreground amino acid; and as phenotype changes, we only considered the acquisition of the foreground phenotype. As coincident changes, we considered the phylogenetic branches where these events coincided. This approach is referred to as “Convergence” (Fig. 1A).

**Fig 1:**
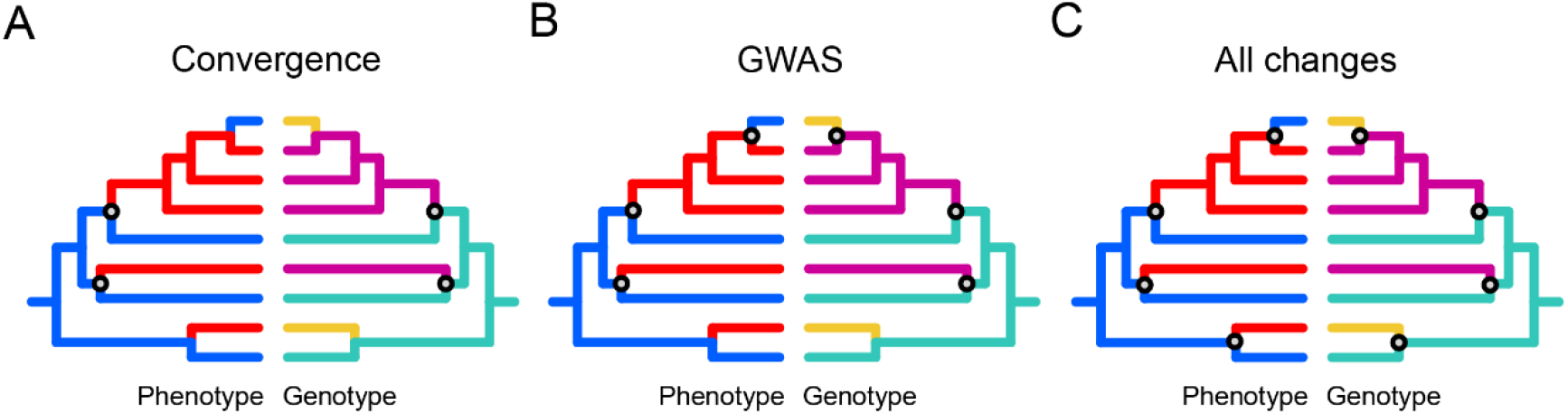
Three approaches to counting the number of simultaneous changes in the phenotype and the encoded amino acid. In each panel, the two trees facing each other show the same phylogeny, with the coloring corresponding to the trait states in the corresponding branches; phenotype in the left, and amino acid at a particular position in the right. Circles indicate changes in phenotype and genotype deemed coincident under the corresponding approach. A, under the Convergence approach, of the three gains of the foreground phenotypic trait state (blue to red), two coincide with the gains of the foreground amino acid (cyan to purple). B, under the GWAS approach, two gains (blue to red) and one loss (red to blue) of the foreground phenotypic trait state coincide respectively with two gains (cyan to purple) and one loss (purple to yellow) of the foreground amino acid. C, under the All changes approach, four changes in the phenotype coincide with four changes in the encoded amino acid. The corresponding numbers of simultaneous events is two (A), three (B) and four (C).

In the second approach, as genotype changes, we considered both the gains and the losses of the foreground amino acid, and as phenotype changes, both the gains and the losses of the foreground phenotype. As coincident changes, we considered the phylogenetic branches where both the foreground amino acid and the foreground phenotype were gained, or both were lost. This approach is referred to as “GWAS” (Fig. 1B).

Finally, in the third approach, we assumed that any amino acid substitution constitutes a genotype change event, and any change in the phenotype counts independently of its direction. This approach is referred to as “All changes” (Fig. 1C).

These approaches correspond to different assumptions regarding the genotype-phenotype association. The “Convergence” approach assumes that the gains of the trait are associated with a gain of a specific amino acid variant, while its losses can proceed through multiple means. The “GWAS” approach assumes that both the gains and the losses of the trait are associated respectively with gains and losses of a specific amino acid variant. Finally, the “All changes” approach assumes that both the gains and the losses of the trait are associated with any changes in the encoded amino acid.

### Change of site-specific amino acid propensities

To get an alternative view of amino acid changes associated with convergent phenotype characteristics, we asked, for each amino acid site, if changes in amino acid propensities correlated with phenotype change. Among many methods for assessment of position-specific amino acid profiles, we chose the Profile Change method (Rey et al, 2018) as it was developed for studies of parallel evolution. This method assigns two amino acid profiles to each site, one for foreground and one for background branches. It then estimates differences between these two profiles in a Bayesian framework and reports the posterior probability that amino acid preferences differ between the two classes of branches. As branch classes, we used the ones for which the corresponding phenotype state was reconstructed.

### 3D Structure

To estimate the functional role of candidate mutations, we reconstructed the 3D structure of genes that carry them. Sequences of *Gallus gallus* genes *ND1, ND2, ND4, ND5* and *ND6* were aligned with homologous genes of *Ovis aries*. The protein 3D structure based on homology with *Ovis aries* respitatory complex I (PDB:5LNK) was reconstructed by Modeller software (Webb and Sali 2016, Marti-Renom et al. 2000, Sali 1993, Fiser et al 2000).

## Results

### Simultaneous change

All three types of simultaneous score metrics revealed that the number of significant associations was low. The highest number of sites (8) was detected in high altitude birds, all of them of marginal significance. The Convergence test detected two sites in *ND1* and *ND5* genes (Figure 2). The GWAS test detected another two sites in *ND2* gene (Figure S1). The All changes test detected 6 sites in *ND1, ND2, ND4, ND5* and *ND6* genes (Figure S3). Findings of different tests partially overlap (Table 2). Additionally, the All changes test detected the site associated with adaptation to long-distance migration in the *ND5* gene. When the stronger Bonferroni correction was applied, only two sites detected by the Convergence test in high altitude birds and one site detected by the All changes test in long-distance migrants remained significant (Table 2).

**Fig 2:**
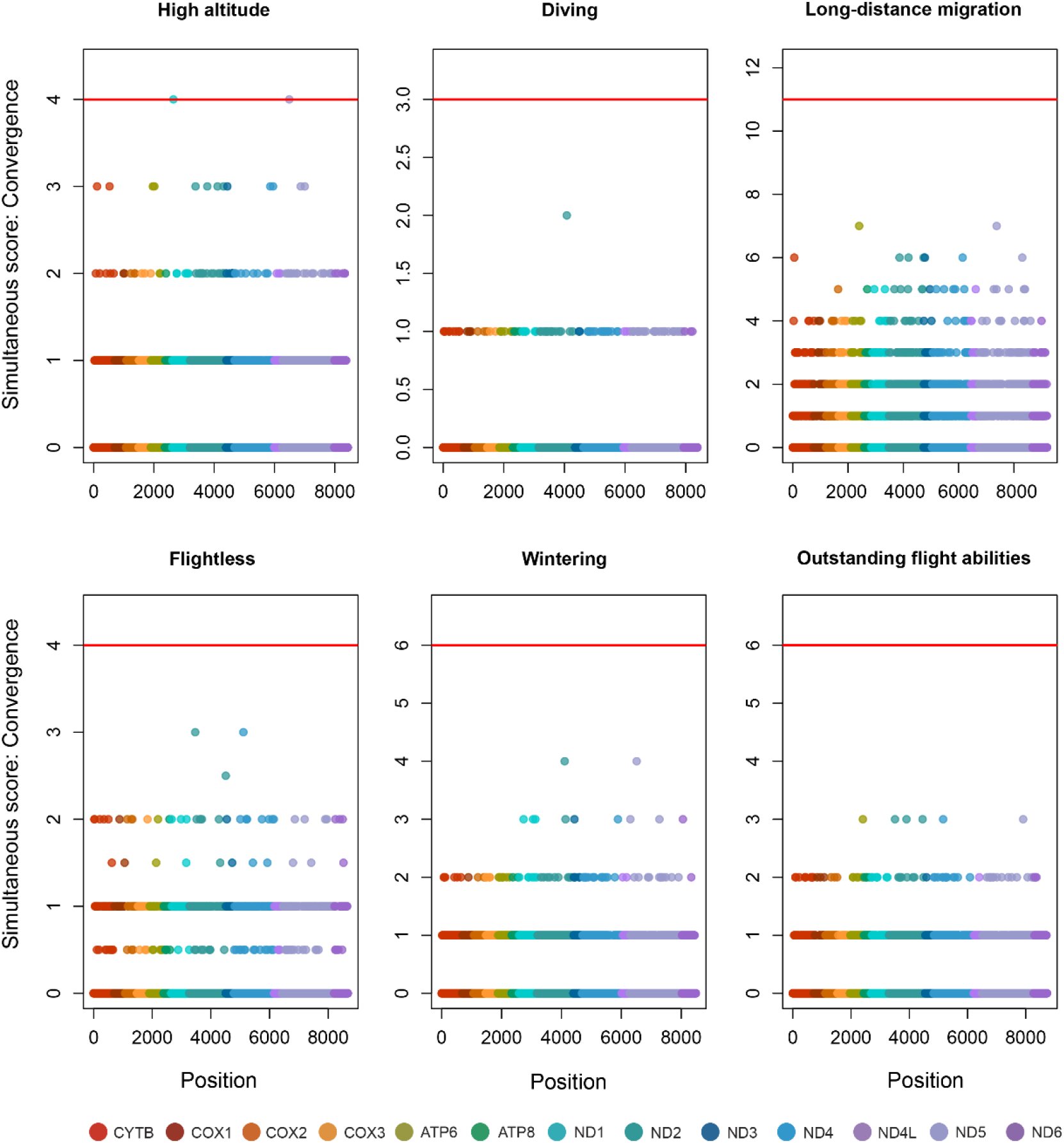
Simultaneous test based on the Convergence approach for each of the six considered phenotypic traits. Horizontal axis, position in the mitochondrial genes; vertical axis, number of simultaneous changes of phenotype and genotype.

**Table 2:** List of significant SNPs detected by the Simultaneous test. Correction 1 column indicates the sites that remain significant after Bonferroni correction accounting for the number of considered sites. Correction 2 column indicates the sites that remain significant after Bonferroni correction accounting for the number of sites and the number of tests (3 tests: Convergence, All changes and GWAS).

| Test type | Gene | Position in *Gallus gallus* | Correction 1 | Correction 2 |
| --- | --- | --- | --- | --- |
| High altitude |  |  |  |  |
| Convergence | ND1 | 17aa(I) | + | + |
|  | ND5 | 57aa(H) | + | + |
| All changes | ND1 | 17aa(I) | + | - |
|  | ND2 | 329aa(T) | + | - |
|  | ND4 | 418(T) | + | - |
|  | ND5 | 114aa(F) | + | - |
|  | ND5 | 495aa(T) | + | - |
|  | ND6 | 135(V) | + | - |
| GWAS | ND2 | 278(M) | + | - |
|  | ND2 | 329aa(T) | + | - |
| Long-distance migration |  |  |  |  |
| All changes | ND5 | 533(T) | + | + |

### Profile change

As an additional test for convergence, we use the amino acid Profile change metric. We expect that recurrent mutations emerging simultaneously with convergent phenotype change could also lead to a change in the amino acid profile between the branches carrying the foreground and the background phenotypes. We compared the results of the Simultaneous tests under all three approaches with the Profile change test (Figure 3, Figure S2, S4).

**Fig. 3:**
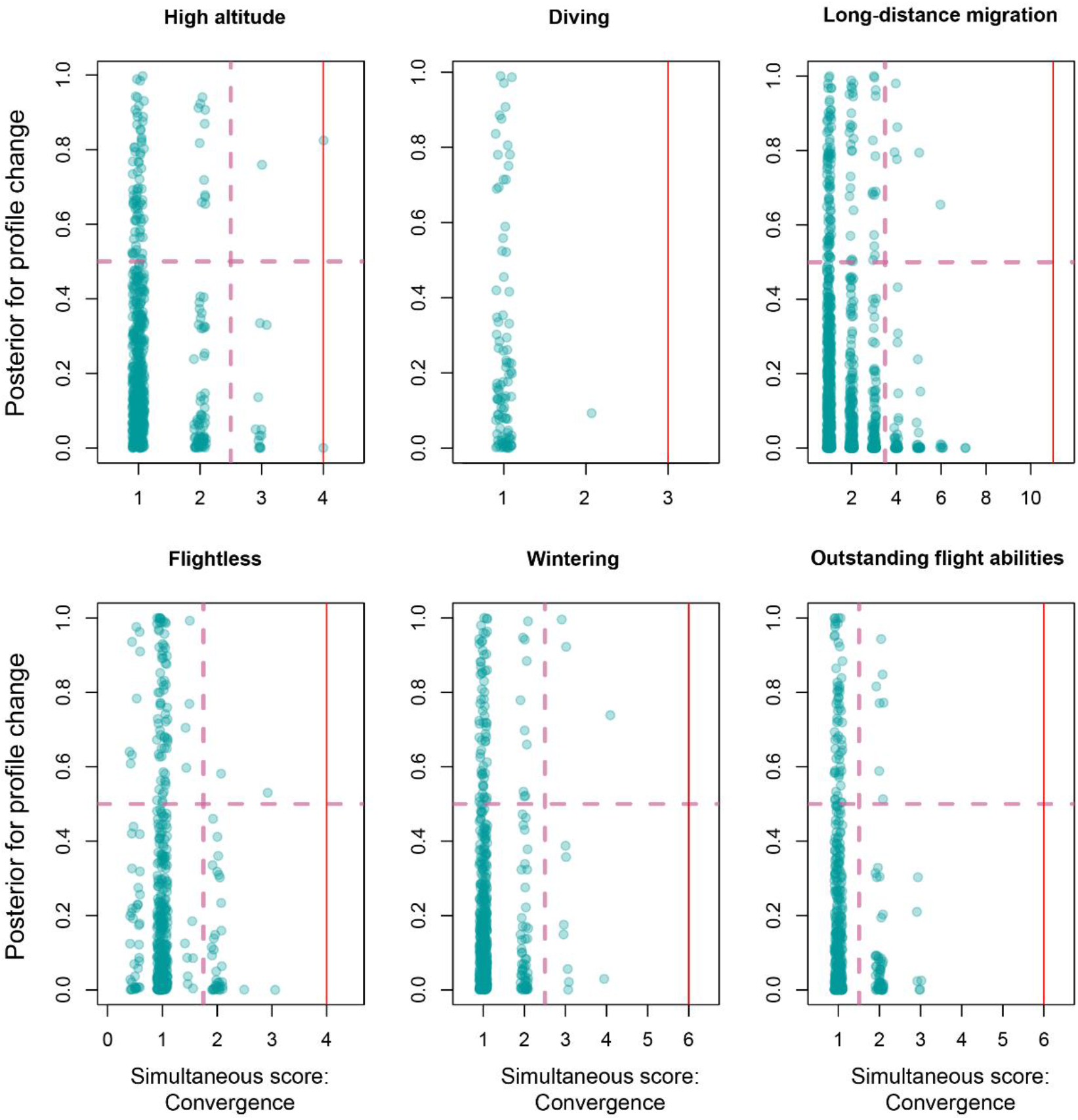
Profile change vs Simultaneous score (Convergence approach). The red line shows the significance threshold for the Simultaneous test. The dashed lines show division of the plot for the Fisher test.

First, we were interested to estimate profile change levels at sites that have a significant Simultaneous score. Among them, only one (57th position of *ND5*, detected by the Convergence test) has profile change score above 0.5 (0.82) and thus can be considered as potentially convergent. Others either result from convergent changes to the same amino acid without a profile change (17th position of *ND1*), or are a consequence of divergent evolution (all other positions).

Second, we could expect that sites with higher simultaneous score could have higher profile change metric if a substantial fraction of these sites is involved in convergent adaptations. To test this assumption, we arbitrarily divided the plots (Figure 3, Figure S2, S4) into four parts, and tested if sites with higher simultaneous score have higher profile change score. We found no dependency in any of the tested phenotype groups (Fisher test, significance level 0.01).

### Sites with evidence for phenotypic association

Position 57 in the *ND5* gene carries the highest signal of functional convergence, as both the Simultaneous and Profile change scores are rather high for it (4 and 0.82 respectively). However, except for the 4 substitutions from Histidine to Tyrosine that occurred synchronously with the phenotype change, there are tens of other substitutions between these two amino acids that occurred at various positions of the phylogeny (partially shown in Figure 4). This suggests that this convergence can be accidental.

**Fig 4:**
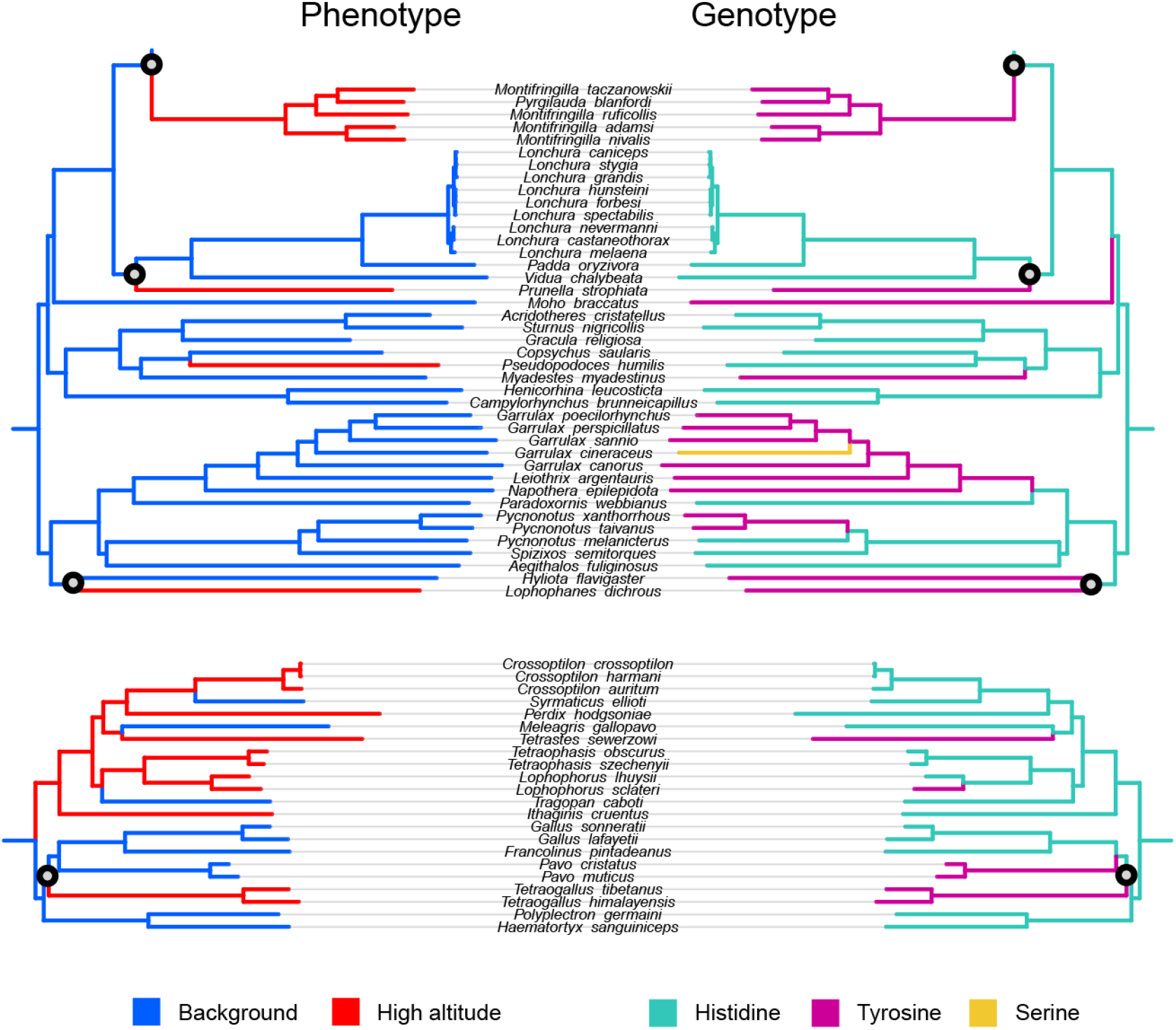
Position 57 in the *ND5* gene associated with high altitude adaptation. This position carries 4 substitutions from Histidine to Tyrosine simultaneous with adaptation, and a Profile change score of 0.82.

Other positions with relatively high Simultaneous scores demonstrate low Profile change scores. Thus, at best some of these sites could be involved in divergent evolution associated with phenotype changes. We run the MEME tool (Kosakovsky Pond et al, 2005) on *ND* genes to find sites involved in recurrent positive selection, yet there was no overlap with our findings.

As all 9 candidate positions were in *ND* genes, we suggested that adaptations could be associated with the respiratory complex I. Among all 3602 analyzed amino acid positions, *ND* genes account for 1998 positions (55%), so it is unlikely to be a coincidence. To ask if there is additional evidence for function, we mapped the candidate positions onto the 3D structure of the respiratory complex I (Figure 5). All the positions are far from the FeS electron-transport clusters and are buried into the membrane arm of the respiratory complex I. They are not grouped together, and they are not close to polar residues in proton channels that play a key role in proton transport (Fiedorczuk, 2016). In total, structural data provides no additional evidence to consider these sites as adaptive.

**Fig 5:**
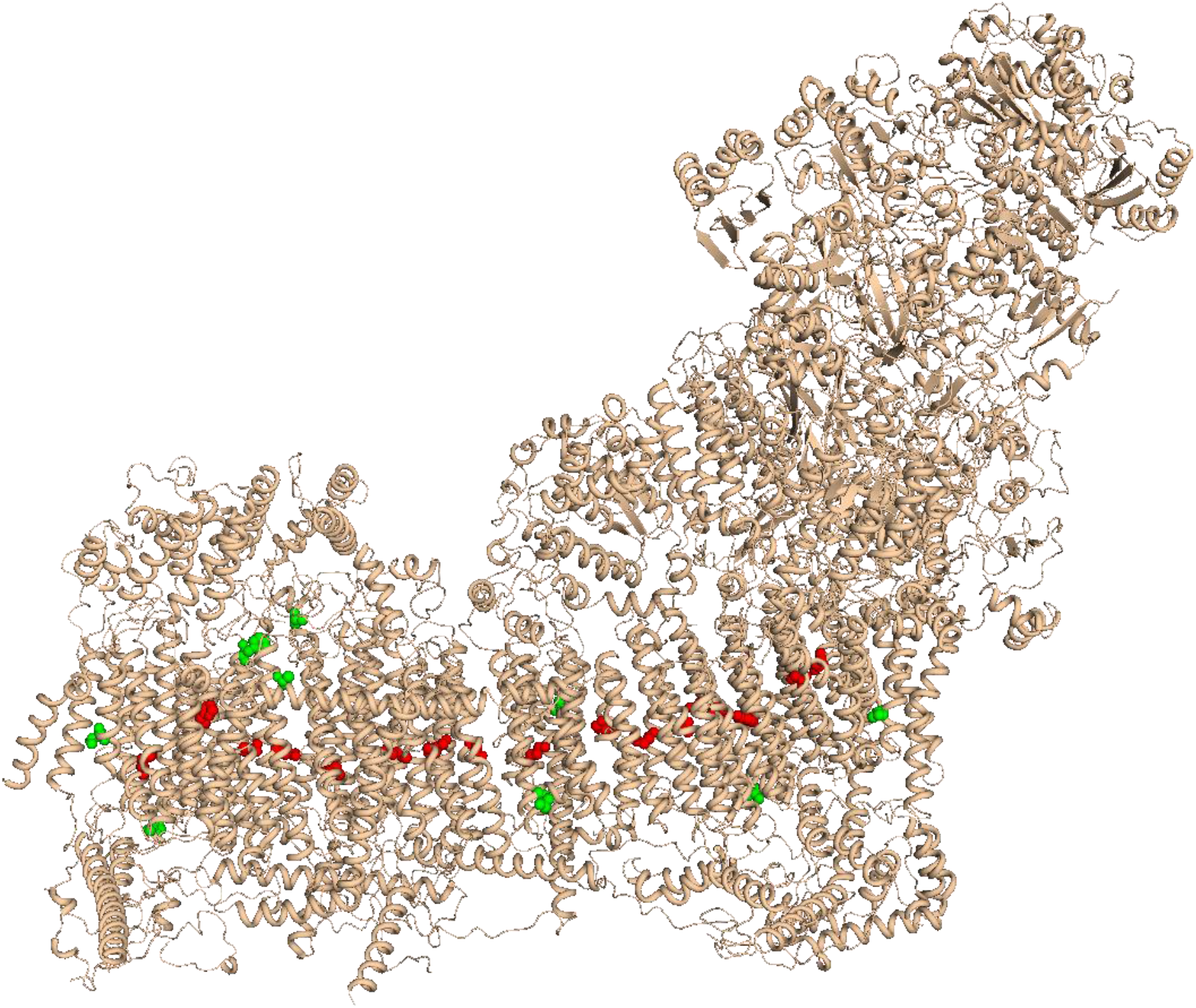
3D structure of respiratory complex I. Candidate amino acid residues are colored green. Polar residues in proton channels, which play a key role in proton transport, are colored red.

## Discussion

The site with the strongest signal of convergence detected in our analysis, which has high Convergence and Profile change scores (position 57 in *ND6*), could originate from functional convergence. Alternatively, the observed pattern of changes could still be coincident, as the mutation pattern rather looks like switches between the set of permitted amino acid under time-invariant amino acid constrains. Among other, basically divergent, positions, none provided additional evidence for selection. We suppose that the concentration of significant associated SNPs in the *ND* genes could be a consequence of higher mutation rates in *ND* genes.

Though we detect no significant associations at the amino acid level, we hypothesized that sites with higher Simultaneous scores could have elevated Profile change scores. This could be the case if instead of a few strong associations, the data carried sites with convergent associations at a small group of phylogenetically close species, or if the associations were weak. However, we detect no excess Profile change score among the sites with higher Simultaneous scores.

Previous works have attempted to detect convergent single-nucleotide mutations in such distant groups as marine mammals (Foote et al, 2015) or echolocating bats and whales (Parker et al, 2013). Many of these attempts failed to find significant convergence or were disproved by later studies (Zou and Zhang 2015, Thomas and Hanh 2015). Similar to those works, we here explore convergence between distantly related species. All the phenotypes analyzed here were acquired repeatedly (Table 1), supposedly making the convergence-based analysis of adaptation more powerful. However, our results did not support this assumption. This suggests that adaptive convergent evolution is rare or hardly detectable in bird OXPHOS system at the considered phylogenetic distances.

There remains a possibility that convergent adaptations in the OXPHOS system could be found in groups of close relatives when the tree of life will be sequenced with higher density. As it was shown by Natarajan et al (2016), similar mutations in hemoglobin subunits lead to high latitude adaptations only in a similar genetic context: there are typical “hummingbird high altitude mutations” and typical “duck high altitude mutations”. If so, the near-lack of signal in our study has to do with the fact that it was based on too distant organisms which have highly divergent evolutionary landscapes in the genes of interest. However, other work indicates that similar substitutions are rarely involved in independent adaptation to high altitudes even inside a group of closely related species like hummingbirds (Lim et al 2019).

Further work may improve the search for simultaneous changes by using better statistical models. Specifically, these models could be improved by incorporating the heterogeneity of the substitution rates between sites or phylogenetic branches.

