## Supplementary_materials for "Convergent adaptation in mitochondria of phylogenetically distant birds: does it exist?"

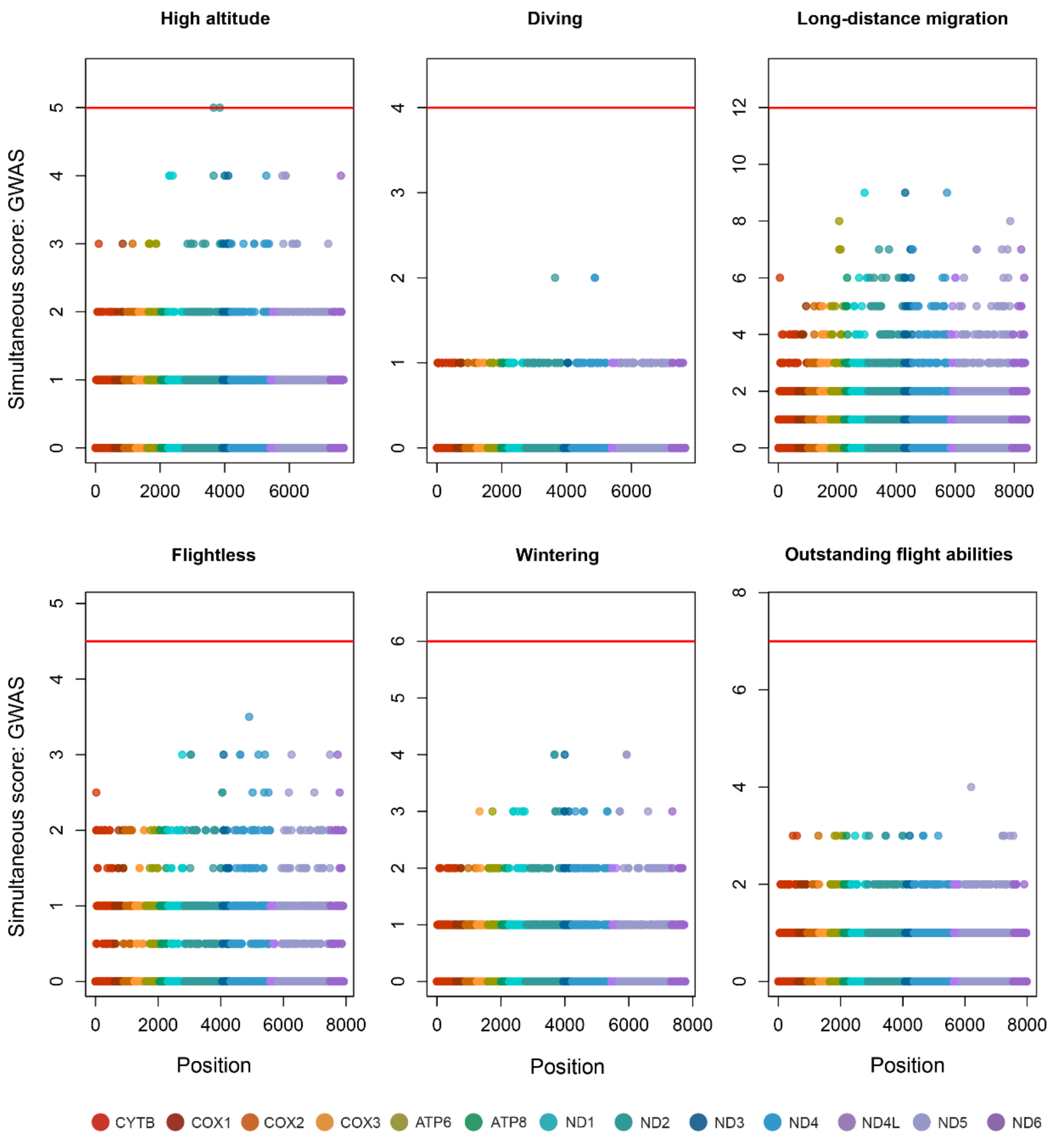


Fig. S1: Simultaneous score. GWAS. Horizontal axis, position in the mitochondrial genes; vertical axis, number of simultaneous changes of phenotype and genotype.

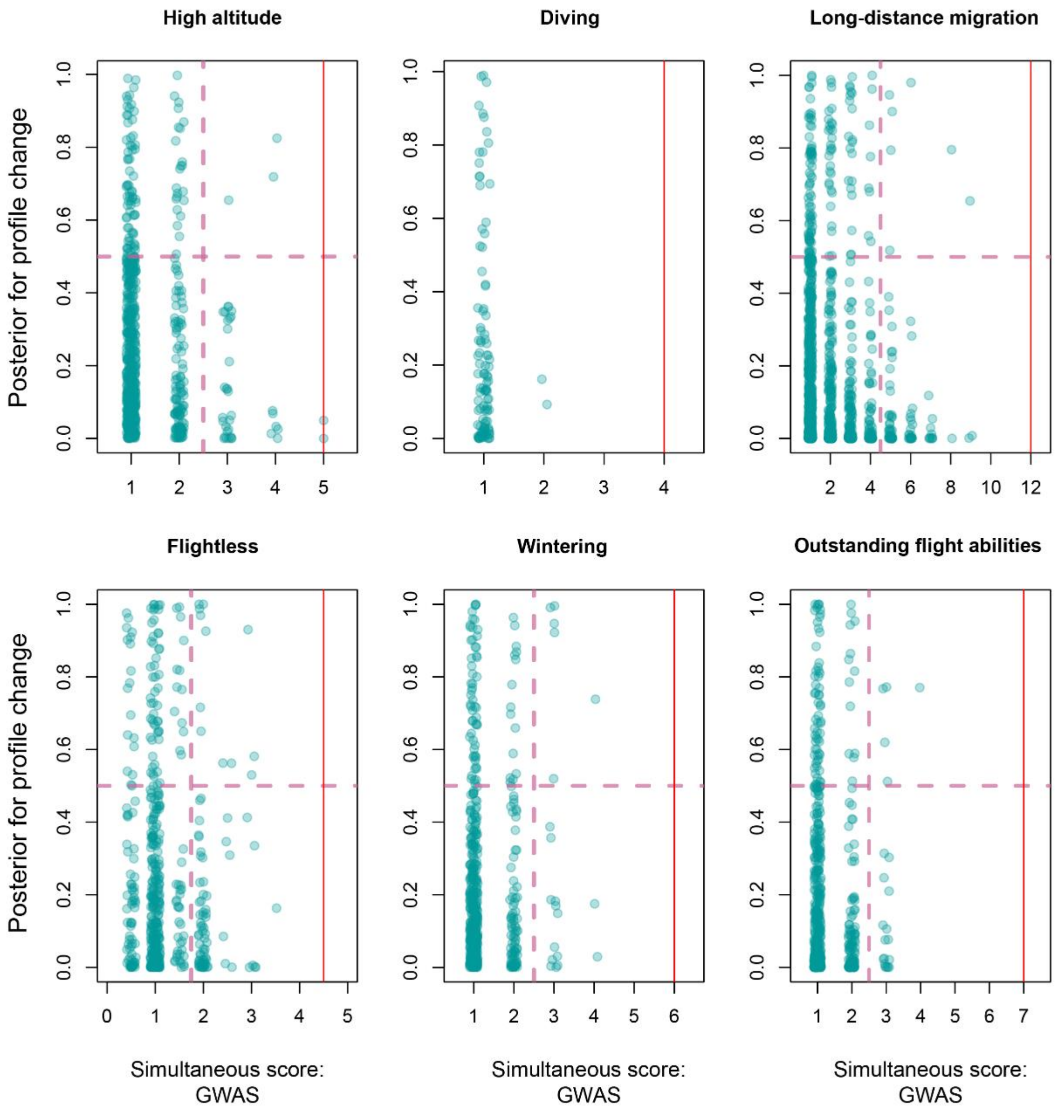


Fig. S2: Profile change vs simultaneous score (GWAS). Red line shows significance threshold for simultaneous test. Dashed lines show division of plot for Fisher test.

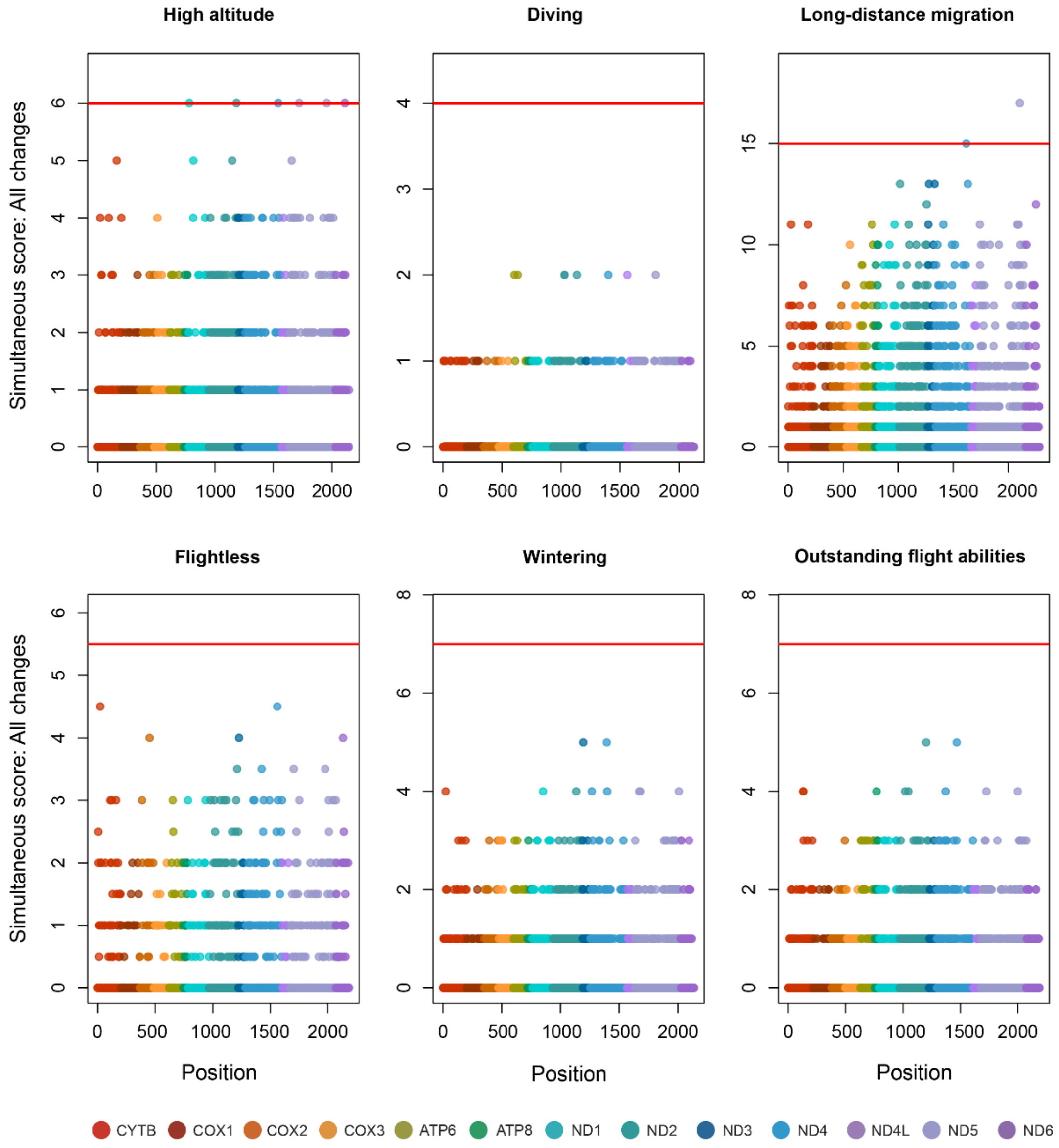


Fig. S3: Simultaneous score. All changes. Horizontal axis, position in the mitochondrial genes; vertical axis, number of simultaneous changes of phenotype and genotype.

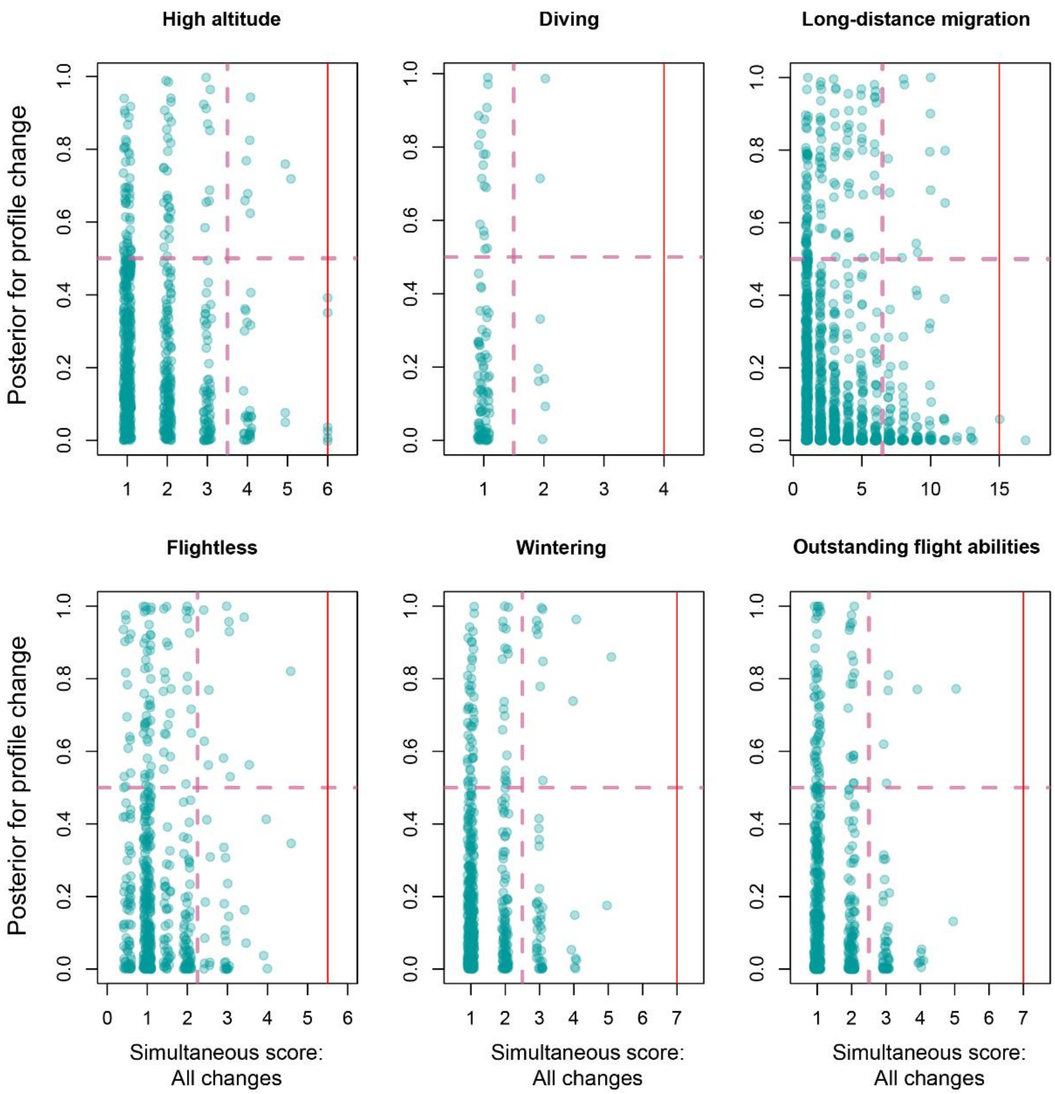


Fig. S4: Profile change vs simultaneous score (All changes). Red line shows significance threshold for simultaneous test. Dashed lines show division of plot for Fisher test.

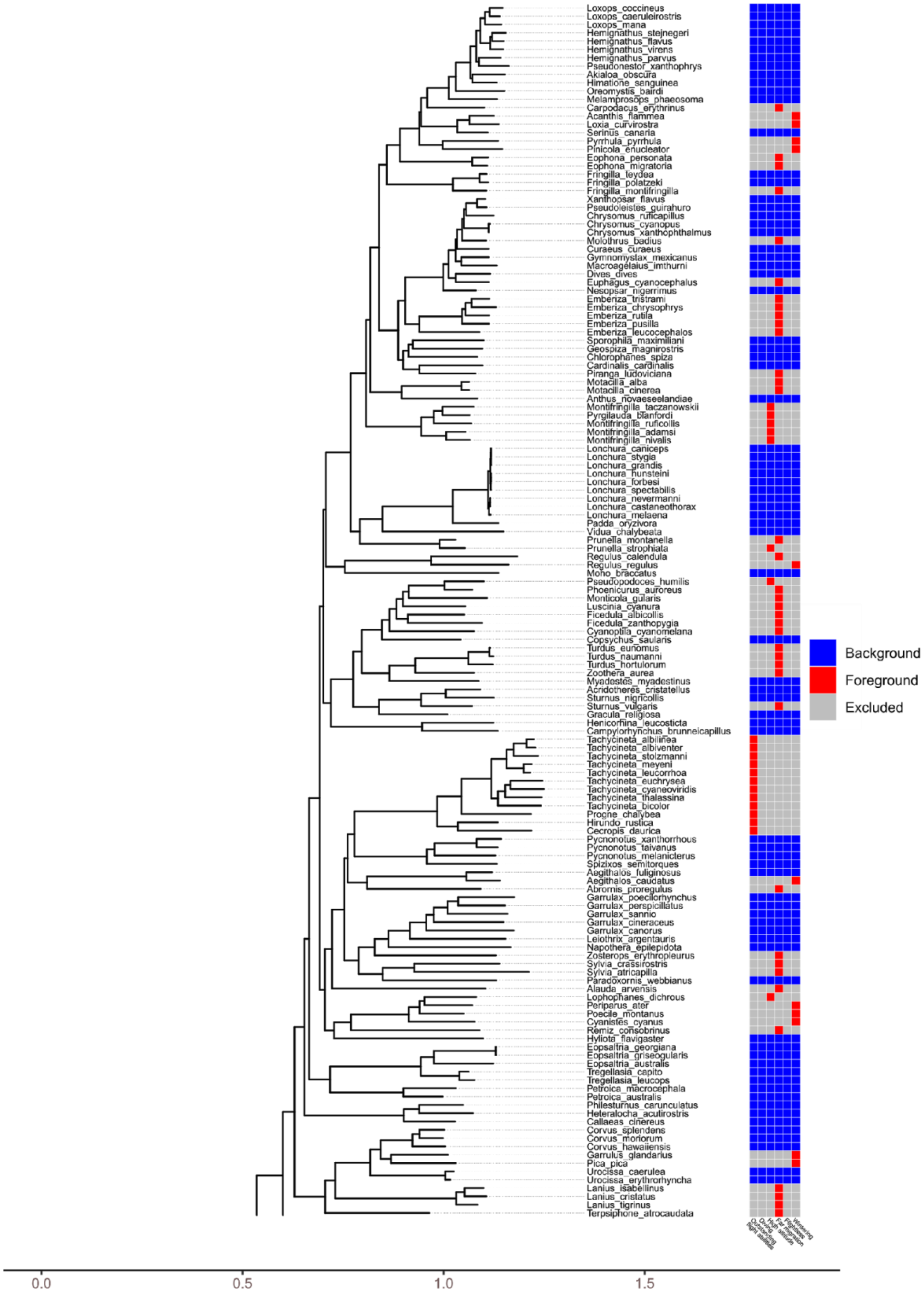
Loxops_coccineus
Loxops_caeruleirostris
Loxops_mana
Hemignathus_stejnegeri
Hemignathus_flavus
Hemignathus_virens
Hemignathus_parvus
Pseudonestor_xanthophrys
Akialoa_obscura
Himatione_sanguinea
Oreomystis_bairdi
Melamprosops_phaeosoma
Carpodacus_erythrinus
Acanthis_flammea
Loxia_curvirostra
Serinus_canaria
Pyrrhula_pyrrhula
Pinicola_enucleator
Eophona_personata
Eophona_migratoria
Fringilla_teydea
Fringilla_polatzeki
Fringilla_montifringilla
Xanthopsar_flavus
Pseudoleistes_guirahuro
Chrysomus_ruficapillus
Chrysomus_cyanopus
Chrysomus_xanthophthalmus
Molothrus_badius
Curaeus_curaeus
Gymnomystax_mexicanus
Macroagelaius_imthurni
Dives_dives
Euphagus_cyanocephalus
Nesopsar_nigerrimus
Emberiza_tristrami
Emberiza_chrysophrys
Emberiza_rutila
Emberiza_pusilla
Emberiza_leucocephalos
Sporophila_maximiliani
Geospiza_magnirostris
Chlorophanes_spiza
Cardinalis_cardinalis
Piranga_ludoviciana
Motacilla_alba
Motacilla_cinerea
Anthus_novaeseelandiae
Montifringilla_taczanowskii
Pyrgilauda_blanfordi
Montifringilla_ruficollis
Montifringilla_adamsi
Montifringilla_nivalis
Lonchura_caniceps
Lonchura_stygia
Lonchura_grandis
Lonchura_hunsteini
Lonchura_forbesi
Lonchura_spectabilis
Lonchura_nevermanni
Lonchura_castaneothorax
Lonchura_melaena
Padda_oryzivora
Vidua_chalybeata
Prunella_montanella
Prunella_strophiata
Regulus_calendula
Regulus_regulus
Moho_braccatus
Pseudopodoces_humilis
Phoenicurus_auroreus
Monticola_gularis
Luscinia_cyanura
Ficedula_albicollis
Ficedula_zanthopygia
Cyanoptila_cyanomelana
Copsychus_saularis
Turdus_eunomus
Turdus_naumanni
Turdus_hortulorum
Zoothera_aurea
Myadestes_myadestinus
Acridotheres_cristatellus
Sturnus_nigricollis
Sturnus_vulgaris
Gracula_religiosa
Henicorhina_leucosticta
Campylorhynchus_brunneicapillus
Tachycineta_albilinea
Tachycineta_albiventer
Tachycineta_stolzmanni
Tachycineta_meyeni
Tachycineta_leucorrhoa
Tachycineta_euchrysea
Tachycineta_cyaneoviridis
Tachycineta_thalassina
Tachycineta_bicolor
Progne_chalybea
Hirundo_rustica
Cecropis_daurica
Pycnonotus_xanthorrhous
Pycnonotus_taivanus
Pycnonotus_melanicterus
Spizixos_semitorques
Aegithalos_fuliginosus
Aegithalos_caudatus
Abrornis_proregulus
Garrulax_poecilorhynchus
Garrulax_perspicillatus
Garrulax_sannio
Garrulax_cineraceus
Garrulax_canorus
Leiothrix_argentauris
Napothera_epilepidota
Zosterops_erythropleurus
Sylvia_crassirostris
Sylvia_atricapilla
Paradoxornis_webbianus
Alauda_arvensis
Lophophanes_dichrous
Periparus_ater
Poecile_montanus
Cyanistes_cyanus
Remiz_consobrinus
Hyliota_flavigaster
Eopsaltria_georgiana
Eopsaltria_griseogularis
Eopsaltria_australis
Tregellasia_capito
Tregellasia_leucops
Petroica_macrocephala
Petroica_australis
Philesturnus_carunculatus
Heteralocha_acutirostris
Callaeas_cinereus
Corvus_splendens
Corvus_moriorum
Corvus_hawaiiensis
Garrulus_glandarius
Pica_pica
Urocissa_caerulea
Urocissa_erythrorhyncha
Lanius_isabellinus
Lanius_cristatus
Lanius_tigrinus
Terpsiphone_atrocaudata
Outstanding flight abilities
Diving
High altitude
Far migration
Flightless
Wintering
Background
Foreground
Excluded
0.0
0.5
1.0
1.5

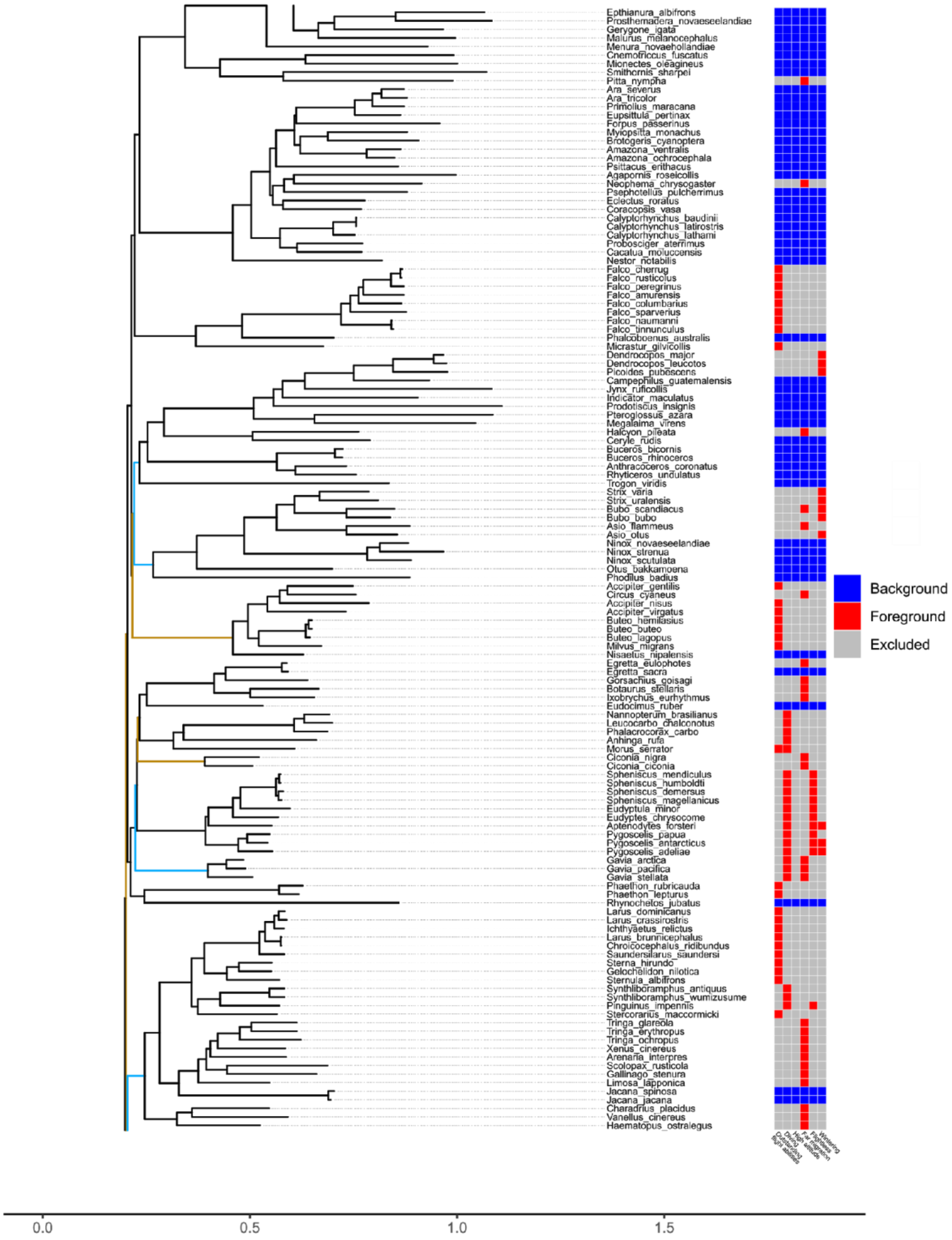

Epthianura_albifrons
Prosthemadera_novaeseelandiae
Gerygone_igata
Malurus_melanocephalus
Menura_novaehollandiae
Cnemotriccus_fuscatus
Mionectes_oleagineus
Smithornis_sharpei
Pitta_nympha
Ara_severus
Ara_tricolor
Primolius_maracana
Eupsittula_pertinax
Forpus_passerinus
Myiopsitta_monachus
Brotogeris_cyanoptera
Amazona_ventralis
Amazona_ochrocephala
Psittacus_erithacus
Agapornis_roseicollis
Neophema_chrysogaster
Psephotellus_pulcherrimus
Eclectus_roratus
Coracopsis_vasa
Calyptorhynchus_baudinii
Calyptorhynchus_latirostris
Calyptorhynchus_lathami
Probosciger_aterrimus
Cacatua_moluccensis
Nestor_notabilis
Falco_cherrug
Falco_rusticolus
Falco_peregrinus
Falco_amurensis
Falco_columbarius
Falco_sparverius
Falco_naumanni
Falco_tinnunculus
Phalcoboenus_australis
Micrastur_gilvicollis
Dendrocopos_major
Dendrocopos_leucotos
Picoides_pubescens
Campephilus_guatemalensis
Jynx_ruficollis
Indicator_maculatus
Prodotiscus_insignis
Pteroglossus_azara
Megalaima_virens
Halcyon_pileata
Ceryle_rudis
Buceros_bicornis
Buceros_rhinoceros
Anthracoceros_coronatus
Rhyticeros_undulatus
Trogon_viridis
Strix_varia
Strix_uralensis
Bubo_scandiacus
Bubo_bubo
Asio_flammeus
Asio_otus
Ninox_novaeseelandiae
Ninox_strenua
Ninox_scutulata
Otus_bakkamoena
Phodilus_badius
Accipiter_gentilis
Circus_cyaneus
Accipiter_nisus
Accipiter_virgatus
Buteo_hemilasius
Buteo_buteo
Buteo_lagopus
Milvus_migrans
Nisaetus_nipalensis
Egretta_eulophotes
Egretta_sacra
Gorsachius_goisagi
Botaurus_stellaris
Ixobrychus_eurhythmus
Eudocimus_ruber
Nannopterum_brasilianus
Leucocarbo_chalconotus
Phalacrocorax_carbo
Anhinga_rufa
Morus_serrator
Ciconia_nigra
Ciconia_ciconia
Spheniscus_mendiculus
Spheniscus_humboldti
Spheniscus_demersus
Spheniscus_magellanicus
Eudyptula_minor
Eudyptes_chrysocome
Aptenodytes_forsteri
Pygoscelis_papua
Pygoscelis_antarcticus
Pygoscelis_adeliae
Gavia_arctica
Gavia_pacifica
Gavia_stellata
Phaethon_rubricauda
Phaethon_lepturus
Rhynochetos_jubatus
Larus_dominicanus
Larus_crassirostris
Ichthyaetus_relictus
Larus_brunnicephalus
Chroicocephalus_ridibundus
Saundersilarus_saundersi
Sterna_hirundo
Gelochelidon_nilotica
Sternula_albifrons
Synthliboramphus_antiquus
Synthliboramphus_wumizusume
Pinguinus_impennis
Stercorarius_maccormicki
Tringa_glareola
Tringa_erythropus
Tringa_ochropus
Xenus_cinereus
Arenaria_interpres
Scolopax_rusticola
Gallinago_stenura
Limosa_lapponica
Jacana_spinosa
Jacana_jacana
Charadrius_placidus
Vanellus_cinereus
Haematopus_ostralegus
Outstanding flight abilities
Diving
High altitude
Far migration
Flightless
Wintering
Background
Foreground
Excluded
0.0
0.5
1.0
1.5

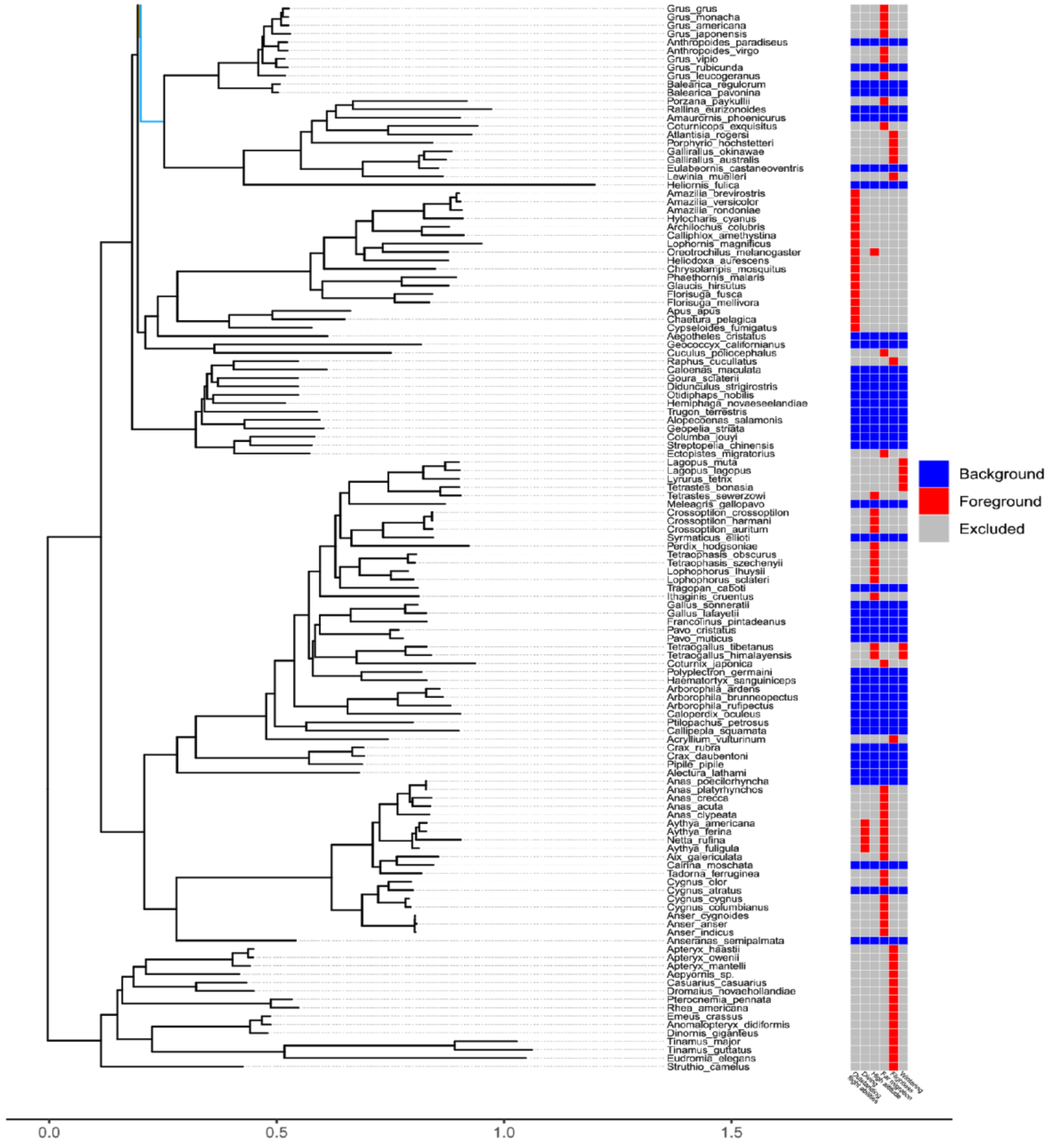


Fig. S5: Phylogentic tree and phenotype groups. Distances are measured in number of substitutions per nucleotide site.
